# Reference-Guided De Novo Retrieval of 3D Chromatin Organizational Features with Representation Learning

**DOI:** 10.64898/2026.10.31.763357

**Authors:** Sean Moran, Jie Liu

**Author notes:** Corresponding author: Jie Liu, Gilbert S. Omenn Department of Computational Medicine & Bioinformatics, University of Michigan, Ann Arbor, MI 48103, USA.

## Abstract

Chromosome conformation maps reveal spatial genome organization through recurring contact patterns, some associated with established architectural features and others with less understood biological processes. Identifying instances of a pattern across these maps is necessary to investigate its biological associations and recurrence across experimental settings. However, conventional feature callers typically target predefined patterns, offering limited support when researchers can provide examples of a feature of interest but lack a dedicated detection method. We introduce Chromatin Fetch (C-Fetch), a reference-guided retrieval framework that uses examples of a desired feature and a contrasting class to rank and assign contact-map patches at provided candidate coordinates. C-Fetch learns a generalized representation through weakly KL-regularized 3D autoencoding of structurally related, non-temporal Hi-C and Micro-C patches without architectural feature labels. We demonstrate retrieval of biologically defined contacts, visually specified patterns, and established architectural features. Polycomb-guided retrieval identifies contacts enriched for PRC2-associated signals, exceeding enrichment from loop-guided assignments at 5 kb, while queries resembling condensed domains retrieve patches with progressively stronger rectangular enrichment at higher ranks. Loop-guided retrieval increases aggregate focal enrichment relative to complete Mustache proposal sets across four held-out maps, and selected contacts show greater focality loss after cohesin depletion than matched remaining candidates. Retrieval also extends to Region Capture Micro-C and HiChIP, with ranking agreement retained across biological replicates and reduced sequencing depths. Together, these results establish reference examples as a flexible means of specifying contact-map features of interest, enabling biological investigation, exploration of visual patterns, and refinement of existing calls across datasets and technologies.

## 1 Introduction

Chromosome conformation capture technologies measure contact frequencies between genomic loci and reveal three-dimensional (3D) chromatin organization (Dekker et al. 2002; Lieberman-Aiden et al. 2009). Hi-C and Micro-C provide genome-wide contact maps, while Region Capture Micro-C and HiChIP examine selected regions or protein-associated interactions (Goel et al. 2023; Mumbach et al. 2016). These maps contain different features such as focal loops, stripes, chromatin jets, and topologically associating domains (TADs) (Rao et al. 2014; Nora et al. 2012; Sexton et al. 2012; Guo et al. 2022; Kim and Kim 2025; Dixon et al. 2012). Most feature-detection methods identify predefined chromatin structures. For example, HiCCUPS and Mustache identify focal contacts (i.e., chromatin loops) using statistical enrichment and peak criteria, whereas other methods detect stripes or domain-like structures (Rao et al. 2014; Roayaei Ardakany et al. 2020; Yoon et al. 2022; Feng et al. 2026). Their scores and thresholds reflect predefined statistical or geometric criteria, offering limited flexibility for follow-up queries that prioritize a different appearance or biological association. Researchers may recognize such patterns through examples before they can formulate a detection rule, even when the patterns lack a precise geometric definition or an established biological interpretation. Existing callers offer limited support for using newly annotated examples to prioritize similar contacts without an additional detection rule or a feature-specific tool. Polycomb-associated chromatin contacts illustrate the need for feature-specific assignment even when a biological association is already known. CTCF/cohesin-associated loops and contacts between Polycomb-associated loci can both appear as focal, long-range enrichments, even though their epigenomic associations can differ (Rao et al. 2014, 2017; Eagen et al. 2017; Kraft et al. 2022; Blackledge and Klose 2021). A loop caller can propose coordinates for both types of contact, but its focal-enrichment score does not establish their Polycomb association. This motivates a shared representation in which reference examples define the contact-map patterns to prioritize.

Building on advances in vision models and representation learning (Caron et al. 2021; He et al. 2022), we investigate whether a shared representation of chromatin conformation contact-map structure can provide a flexible, robust basis for identifying and exploring diverse patterns of chromatin organization. We introduce Chromatin Fetch (C-FETCH), a representation learning framework, for retrieving contact-map features specified through reference examples. At its core, C-Fetch uses a weakly KL-regularized 3D autoencoder to learn representations of groups of structurally related, non-temporal contact-map patches as clips through reconstruction without architectural feature labels (Bengio et al. 2013; Hinton and Salakhutdinov 2006; Kingma and Welling 2014; Rombach et al. 2022; WanTeam et al. 2025). To use C-Fetch, a researcher supplies a chromatin contact map and candidate coordinates, then defines the retrieval target using annotated reference examples and a contrasting negative reference set. Candidate coordinates are genomic coordinate pairs that specify where C-Fetch extracts contact-map patches for feature-class assignment. Reference examples are annotated patches showing the feature of interest and contrasting negative examples; they guide C-Fetch’s rankings and feature assignments. We provide loop and Polycomb-associated reference sets with accompanying negative examples, and make it possible for researchers to define other retrieval targets.

We show that C-Fetch’s representation behaves optimally for reference-guided retrieval, supported by its label-consistent neighborhoods, correspondence between latent proximity and visual resemblance, and improved consistency of similarity rankings when averaging across patch-clip contexts. We then demonstrate three applications of this representation: first, assigning Polycomb-associated contacts as an example of biologically defined feature that conventional loop callers do not specifically identify; second, retrieving a contact-map pattern defined entirely by its visual appearance; and third, we demonstrate improved focal-contact prioritization in loop retrieval over the baseline. Using a set of standardized candidate coordinates, Polycomb-guided C-Fetch assignments showed PRC2-associated enrichment, with stronger signal under Polycomb than loop guidance at 5 kb. This supports example-guided assignment of biologically relevant contact populations within an existing caller’s proposals. For the entirely visually defined query, examples of rectangular enrichment resembling reported condensed-domain contact-map patterns (Paggi et al. 2026) provided a starting point for retrieving and investigating similar patches across chromatin contact map datasets. Rankings tracked the strength of the requested appearance under C-Fetch’s viewing normalization, providing a retrieved population for exploratory analysis of compartment and accessibility characteristics. For chromatin loop-guided retrieval, C-Fetch selects contacts with greater aggregate focal enrichment than the complete Mustache proposal sets. Expanding the candidate pool extends retrieval while preserving most earlier selections and substantial ranking agreement. Selected contacts also show greater focality loss after cohesin depletion than matched remaining candidates, supporting preferential retrieval of contacts with cohesin-sensitive focal structure.

Finally, we demonstrate our reference-guided retrieval transfers to Region Capture Micro-C and HiChIP contact map technologies, that candidate coordinate rankings retain useful agreement across biological replicates and under moderate reductions in sequencing depth, and show by ablating the size of the reference set that larger reference collections improve retrieval consistency and smaller set sizes still support exploratory retrieval of focally enriched populations. Together, the applications show how researchers can use our learned contact-map representation to retrieve biologically defined features, investigate visually specified patterns, and improve selection within established callers.

## 2 Results

### 2.1 Chromatin-Fetch learns a shared representation for reference-guided feature retrieval

Chromatin-Fetch (C-Fetch) learns to represent contact-map patches without architectural feature labels, and then uses selected examples to guide retrieval after training. We trained 3D autoencoders to reconstruct unlabeled Hi-C and Micro-C submatrices (Figure 1A). Model inputs were constructed from raw contact values using C-Fetch’s patch-normalization procedure, without KR matrix balancing (Methods, Section 4.2). Patches with broadly similar contact-map structure were then grouped into clips, which the 3D autoencoder encoded jointly (Methods, Section 4.3). After training, the encoder was frozen, and manually annotated reference examples defined the feature queries used to rank and select patches at supplied candidate coordinates (Figure 1B; Methods, Section 4.5).

**Figure 1:**
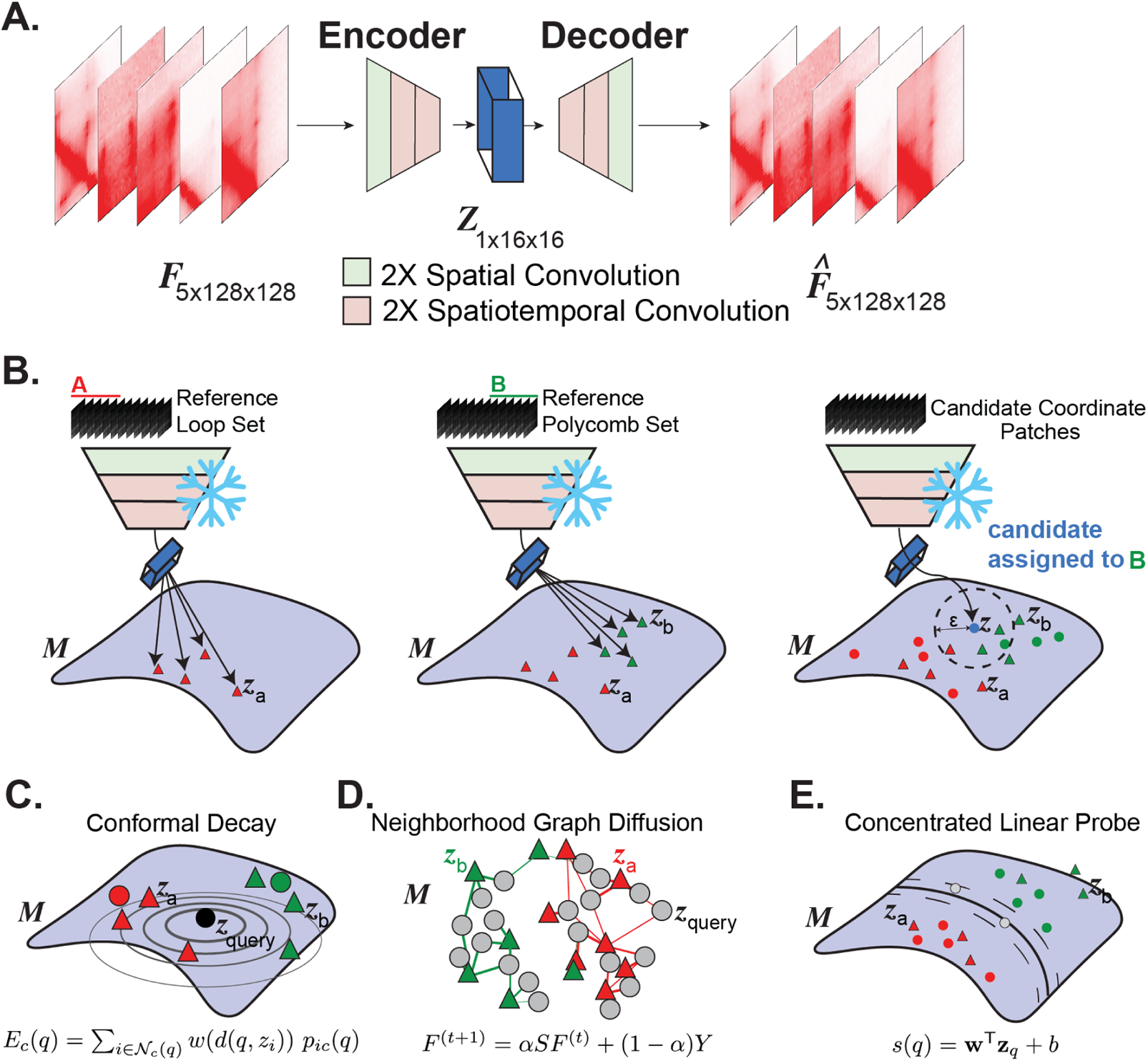
Overview of C-Fetch representation learning and reference-guided inference. (A) Five structurally related Hi-C or Micro-C patches are randomly selected from each eight-frame flatpack record to form a non-temporal input to the weakly KL-regularized 3D autoencoder. The model is trained to reconstruct these inputs without architectural feature labels. (B) After training, the decoder is removed and the frozen encoder maps reference examples and target candidates into a shared continuous representation. The same representation supports different post-training queries by changing the reference examples rather than retraining the encoder. (C) Distance-decayed conformal inference evaluates each candidate using calibrated evidence from nearby reference examples, with the contribution of each example decreasing with latent distance. (D) Graph-diffusion inference propagates reference evidence through a local latent-neighborhood graph and assigns candidates according to the accumulated evidence. (E) Calibrated linear-probe inference fits a linear decision function to frozen reference embeddings and calibrates its candidate assignments using held-out reference examples. Together, these complementary routes convert the frozen C-Fetch representation into reference-guided feature assignments.

For retrieval, a researcher supplies candidate contact-map patches together with reference examples representing the feature of interest and a contrasting class, using either our provided reference sets or independently annotated examples. C-Fetch forms clips from examples within each reference class and places each candidate in several clips with structurally related candidates. The frozen 3D encoder maps these clips into the same latent representation, where conformal scoring, graph diffusion, and the calibrated linear probe evaluate each candidate against the references (Figure 1B–E). C-Fetch returns procedure-specific scores and assignments, a combined ranking, and a consensus assignment for the supplied candidates. Researchers can change the retrieval target by replacing the reference examples without retraining the encoder. To demonstrate this workflow, we evaluate C-Fetch’s retrieval performance on a Polycomb-associated feature set, a visually defined condensed-domain set, and loop queries within Mustache-proposed candidate pools, and differing sequencing depths and contact-map technologies.

The 3D model jointly encoded short, non-temporal clips of structurally related patches, allowing each representation to incorporate local context without implying a biological sequence (Methods, Section 4.3). Their joint encoding supplied non-temporal context without imposing a biological sequence. Both models retained a Gaussian posterior and a small KL penalty, but reconstruction dominated the training objective. Consistent with continuous autoencoders used in latent image and video models, we describe them operationally as weakly KL-regularized autoencoders rather than assuming a calibrated generative prior (Rombach et al. 2022; WanTeam et al. 2025). Unless otherwise specified, subsequent C-Fetch assignment and epigenomic analyses use the base 650M-parameter 3D checkpoint. We evaluate its neighborhood organization and the alternative checkpoints below.

### 2.2 The continuous representation supports similarity-based ranking across datasets

Reference-guided retrieval requires a representation in which proximity reflects relevant contact-map similarities. We evaluated this requirement through agreement with annotated feature labels, resemblance between neighboring patches, and consistency of similarity rankings across sampled contact-map clips.

We first evaluated label agreement by measuring how often an example’s nearest neighbors shared its annotated feature label. We defined neighbors using RMS-L2 distance between latent vectors and calculated neighborhood purity as the mean same-label fraction among the *k* (*k*=25) nearest neighbors. Each of 12 dataset-line–resolution conditions contained *n_p_* (*n_p_*=400) annotated strong-positive and *n_n_* (*n_n_*=400) negative examples, giving an approximately 50% chance baseline. We observed that both base models formed neighborhoods with label agreement substantially above the approximately 50% chance baseline. Increasing model capacity further improved this organization: mean neighborhood purity rose from 0.781 for the 135M model to 0.906 for the 650M model, with higher purity in every evaluated condition (Figure 2A,B; Supplemental Table S1). We also evaluated label agreement when examples from different datasets and resolutions shared the same search population. Purity remained high, reaching 0.906 for the 135M model and 0.973 for the 650M model across all 9,600 examples, with the larger model also performing better when examples were pooled separately by resolution or dataset (Table 1). The larger model maintained higher purity as neighborhood coverage expanded (Figure 2C). The larger model’s advantage therefore extended beyond queries with the most readily populated local neighborhoods. Local decoder-Jacobian spectra decayed smoothly across the 16 measured directions for both models (Figure 2D). Capacity-related gains were also observed under standardized cosine similarity, but were less consistent than under RMS-L2, with purity decreasing in the pooled 10 kb subset (Table 1).

**Figure 2:**
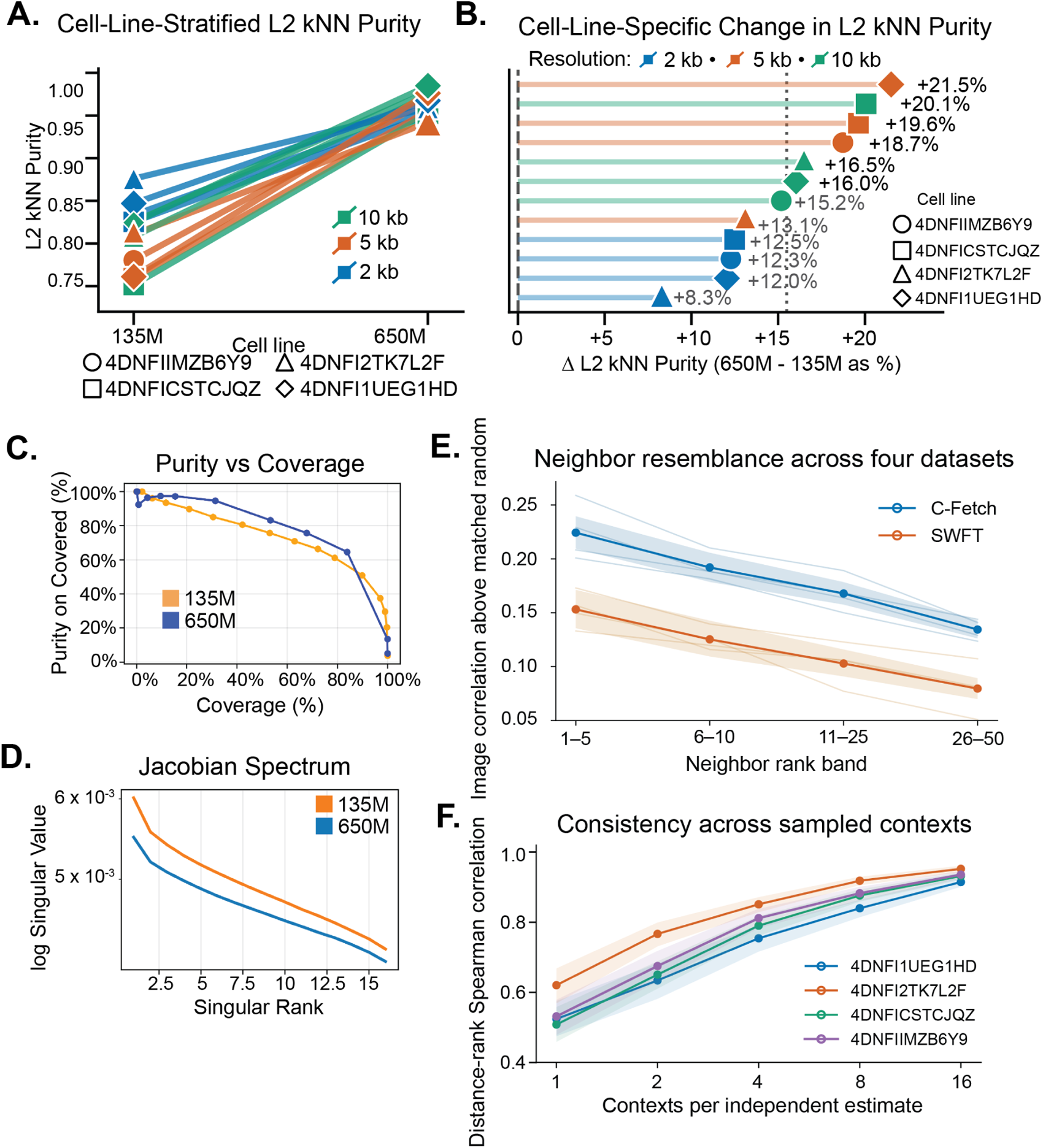
The C-Fetch representation supports similarity-based neighborhood ranking. (A) *k* = 25 RMS-L2 neighborhood purity for the base 135M and 650M 3D models across 12 dataset–resolution conditions, each containing 400 strong-positive and 400 negative inputs. (B) Within-condition differences, Δ purity = 650M *−* 135M. (C) Neighborhood purity versus coverage as the RMS-L2 radius increases. (D) Mean local decoder-Jacobian singular-value spectra estimated using 16 orthonormal latent directions. (E) Image correlation above matched random comparisons across neighbor-rank bands for C-Fetch and SWFT in four datasets at 5 kb. Bold curves show cross-target means; faint curves show individual targets. (F) Spearman agreement between candidate-distance rankings across independently sampled clip contexts. Shading in (E,F) indicates 95% bootstrap intervals; evaluation and resampling procedures are described in Supplemental Methods.

**Table 1:**
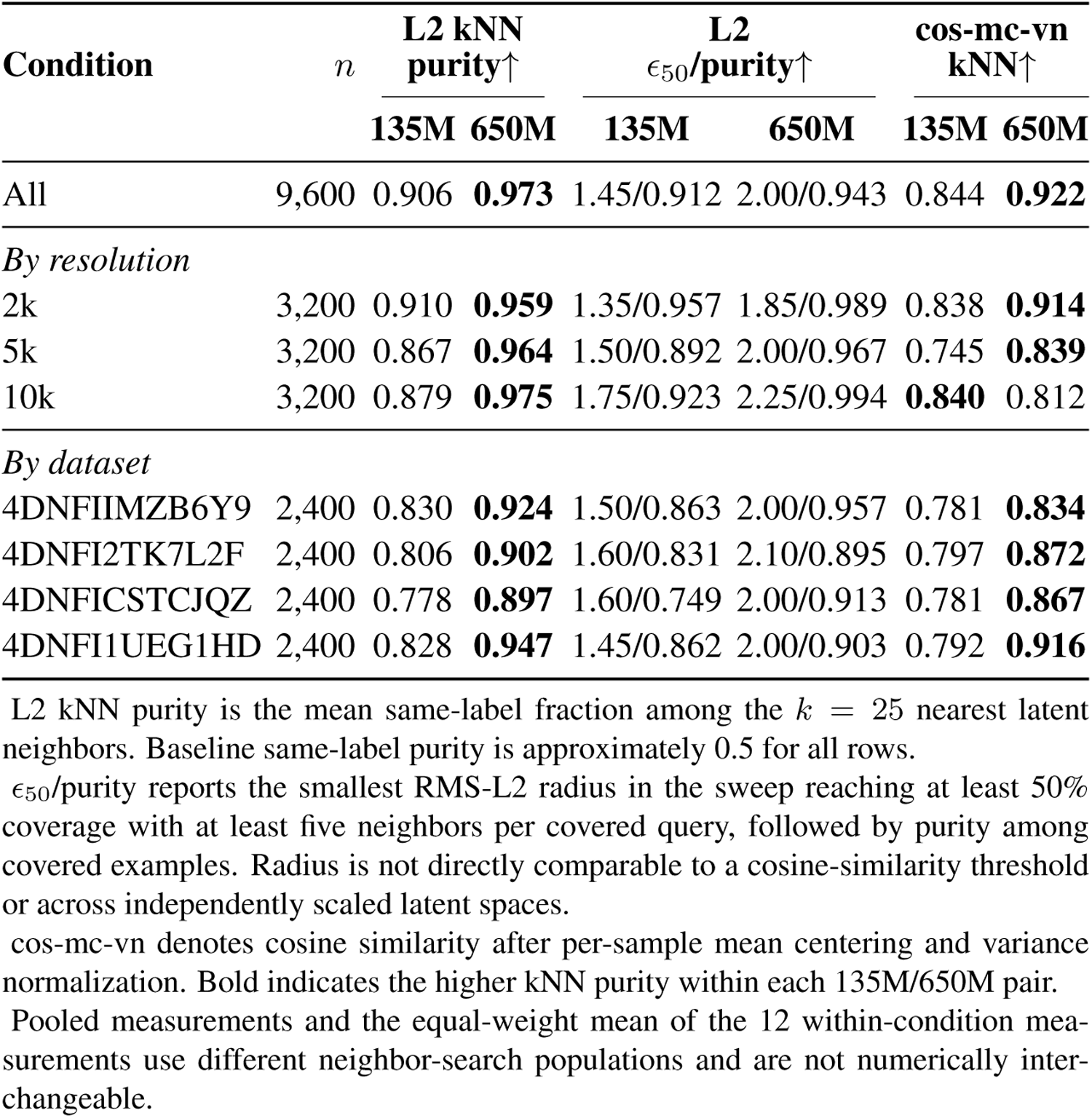
Latent-neighborhood geometry of the 3D autoencoders across accumulated evaluation conditions.

| **Condition** | *n* | **L2 kNN purity↑** |  | **L2 ϵ<sub>50</sub>/purity↑** |  | **cos-mc-vn kNN↑** |  |
|---|---|---|---|---|---|---|---|
|  |  | **135M** | **650M** | **135M** | **650M** | **135M** | **650M** |
| All | 9,600 | 0.906 | **0.973** | 1.45/0.912 | 2.00/0.943 | 0.844 | **0.922** |
| *By resolution* |  |  |  |  |  |  |  |
| 2k | 3,200 | 0.910 | **0.959** | 1.35/0.957 | 1.85/0.989 | 0.838 | **0.914** |
| 5k | 3,200 | 0.867 | **0.964** | 1.50/0.892 | 2.00/0.967 | 0.745 | **0.839** |
| 10k | 3,200 | 0.879 | **0.975** | 1.75/0.923 | 2.25/0.994 | **0.840** | 0.812 |
| *By dataset* |  |  |  |  |  |  |  |
| 4DNFIIMZB6Y9 | 2,400 | 0.830 | **0.924** | 1.50/0.863 | 2.00/0.957 | 0.781 | **0.834** |
| 4DNFI2TK7L2F | 2,400 | 0.806 | **0.902** | 1.60/0.831 | 2.10/0.895 | 0.797 | **0.872** |
| 4DNFICSTCJQZ | 2,400 | 0.778 | **0.897** | 1.60/0.749 | 2.00/0.913 | 0.781 | **0.867** |
| 4DNFI1UEG1HD | 2,400 | 0.828 | **0.947** | 1.45/0.862 | 2.00/0.903 | 0.792 | **0.916** |

L2 kNN purity is the mean same-label fraction among the *k* = 25 nearest latent neighbors. Baseline same-label purity is approximately 0.5 for all rows.
ϵ<sub>50</sub>/purity reports the smallest RMS-L2 radius in the sweep reaching at least 50% coverage with at least five neighbors per covered query, followed by purity among covered examples. Radius is not directly comparable to a cosine-similarity threshold or across independently scaled latent spaces.
cos-mc-vn denotes cosine similarity after per-sample mean centering and variance normalization. Bold indicates the higher kNN purity within each 135M/650M pair.
Pooled measurements and the equal-weight mean of the 12 within-condition mea-surements use different neighbor-search populations and are not numerically inter-changeable.

As an additional test beyond label agreement, we evaluated whether patches closer together in the C-Fetch representation were more visually similar. Using the base 650M model, we ranked candidate neighbors by mean latent distance across sampled clip contexts for 128 query patches in each of four datasets at 5 kb. We measured pixel resemblance relative to randomly sampled comparison patches matched for genomic separation and contact-density characteristics, and compared the results with a fixed windowed Fourier-feature (SWFT) baseline. Nearer C-Fetch neighbors showed greater excess resemblance, and C-Fetch’s mean excess resemblance exceeded that of SWFT in every displayed rank band (Figure 2E).

Finally, because C-Fetch uses related patch clips for embedding, we tested whether permuting clips contents altered its ranking of a specific candidate patch’s neighbors. For each query, we compared rankings obtained from two separately sampled sets of clip contexts while holding the comparison bank fixed. Across the four datasets, mean Spearman agreement increased from 0.508–0.621 with one context per estimate to 0.915–0.952 with 16 contexts, showing that averaging across contexts made similarity rankings more reproducible (Figure 2F). The overlap between the two sets of 25 nearest neighbors also increased with context count, although their membership remained only partially overlapping.

Together, these analyses showed that increased model capacity improved label-consistent neighborhoods, latent proximity tracked patch resemblance, and averaging across clip contexts improved the reproducibility of similarity rankings. These findings support using the representation to compare reference examples with candidate patches; the following application analyses test whether those comparisons guide useful retrieval.

### 2.3 Polycomb guidance shifts an overlapping focal-contact population toward greater PRC2 enrichment

We applied the learned C-Fetch representation to retrieve Polycomb-associated contacts within a broader set of focal-contact candidates. Mustache can propose both Polycomb-associated contacts and CTCF/cohesin-associated loops, which can share a punctate appearance. We asked whether Polycomb-associated reference examples could prioritize contacts with stronger PRC2-associated signal within this candidate pool. We compared Polycomb- and loop-guided selections, evaluated their H3K27me3, EZH2, and SUZ12 enrichment, and examined whether contacts unique to each query differed in these associations.

We used Polycomb-associated reference examples to guide C-Fetch ranking and selection within Mustache-proposed candidate coordinates, and bench-marked retrievals using loop reference examples. The encoder, candidate coordinates, and assignment procedure were held fixed between the two retrievals. For each target dataset, references originating from that dataset were excluded, so retrieval was guided by examples from the other three reference datasets (Figure 3A). We evaluated the resulting selections across four target datasets at 2 kb, 5 kb, and 10 kb resolution. Polycomb and loop references selected overlapping but distinct subsets of the Mustache candidates. At 5 kb, 54.4–81.8% of Polycomb-assigned contacts were also selected using loop references, while each query retained contacts absent from the other selection (Figure 3B; Supplemental Table S2). Thus, reference choice differentiated candidate membership despite substantial overlap between the retrieved populations.

**Figure 3:**
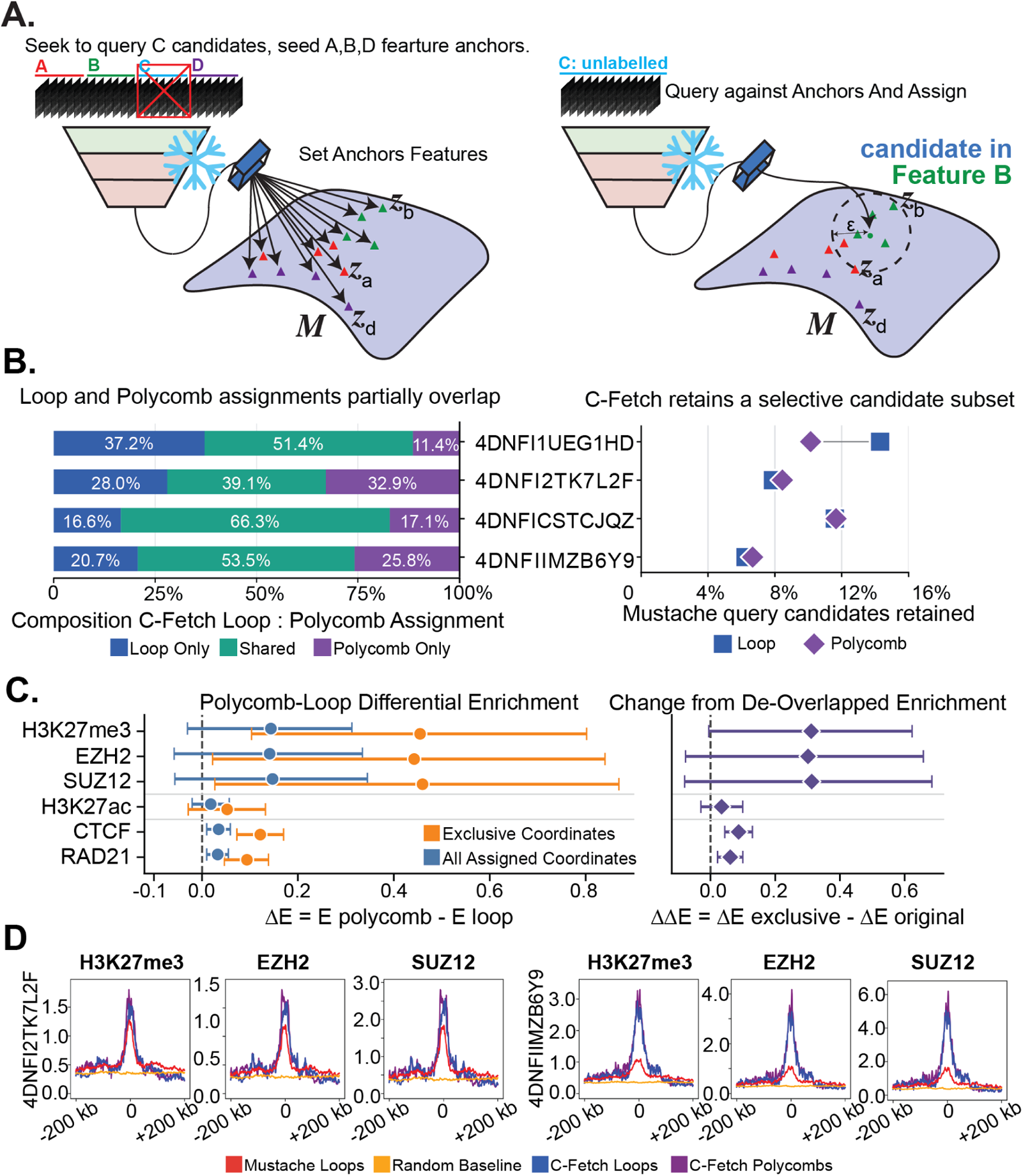
Changing the reference set redirects C-Fetch retrieval. (A) Dataset-held-out comparison using loop or Poly-comb reference examples from the remaining datasets, with the encoder, Mustache candidate coordinates, and assignment procedure held fixed. (B) Selection overlap at 5 kb. Stacked bars partition the selected-coordinate union into loop-only, shared, and Polycomb-only contacts; paired symbols show the percentage of Mustache candidates retained under each query. (C) Polycomb–loop differential enrichment before and after removing shared coordinates, averaged across four target datasets using *±*25-kb anchor windows. Bars show joint-bootstrap 95% confidence intervals. The aligned panel shows the paired change, ΔΔ*E* = Δ*E*_exclusive_ *−* Δ*E*_complete_. (D) Representative H3K27me3, EZH2, and SUZ12 pileups for H1-hESC Micro-C (4DNFI2TK7L2F) and Hi-C (4DNFIIMZB6Y9), comparing Mustache candidates, random-background coordinates, and loop- and Polycomb-guided selections. Signal magnitudes are interpreted within assays, not across marks. Target-level estimates are provided in Supplemental Table S3.

To assess the biological associations of the retrieved contacts, we examined H3K27me3 and the PRC2 components EZH2 and SUZ12, which provide complementary evidence of Polycomb-associated chromatin (Cao et al. 2002; Pasini et al. 2004; Blackledge and Klose 2021). At 5 kb, both retrieved contact populations were strongly enriched relative to random coordinates. Within *±*25-kb windows centered on each contact anchor, median enrichment across targets was 3.11-fold versus 2.71-fold for H3K27me3, 3.89-fold versus 3.36-fold for EZH2, and 4.48-fold versus 3.84-fold for SUZ12 in Polycomb- and loop-guided selections, respectively (Supplemental Table S3). Poly-comb guidance strengthened the enrichment of biologically related polycomb factors relative to loop guidance.

At 5 kb, Polycomb-guided selections showed greater mean PRC2-associated signal within *±*25 kb of the contact anchors. Differential log_2_ enrichment (Δ*E* = *E*_Polycomb_ *−E*_Loop_), averaged equally across the four targets, was 0.141 for H3K27me3, 0.140 for EZH2, and 0.147 for SUZ12, whereas H3K27ac remained near parity (Figure 3C; Supplemental Table S4). Estimates were positive in all four targets for each PRC2-associated measurement, although individual-target confidence intervals included zero except for H3K27me3 in 4DNFI2TK7L2F (Supplemental Figure S1A; Supplemental Table S3). Representative profiles from 4DNFI2TK7L2F and H1-hESC/4DNFIIMZB6Y9 illustrate the shared central PRC2-associated enrichment and generally stronger signal under Polycomb guidance (Figure 3D).

Because many contacts were shared between the selections, we next asked whether contacts unique to the Polycomb query also showed stronger PRC2-associated enrichment than those unique to the loop query. After removing shared coordinates, mean 5 kb Polycomb-minus-loop differential enrichment was 0.452 for H3K27me3, 0.451 for EZH2, and 0.472 for SUZ12, with joint-bootstrap confidence intervals excluding zero for all three measurements (Figure 3C; Supplemental Figure S1B; Supplemental Table S4). These differences were numerically larger than those for the complete selections. The paired increase for H3K27me3 was 0.312, with a 95% confidence interval excluding zero (0.006–0.607), whereas paired confidence intervals for the increases in EZH2 and SUZ12 included zero (Supplemental Table S4). Both exclusive populations nevertheless remained PRC2-enriched relative to matched controls, indicating a relative preference rather than separation into distinct biological classes (Supplemental Figure S1C; Supplemental Table S3). This preference also extended to continuous scores: higher Polycomb-minus-loop reference-preference scores were modestly associated with greater PRC2-associated anchor signal in both H1 datasets (Spearman *ρ* = 0.109–0.169; Supplemental Figure S2). This reference-dependent preference varied with genomic resolution and was most coherent at 5 kb (Supplemental Figure S3; Supplemental Table S3). At 2 kb, de-overlap frequently produced positive PRC2-associated differences, but their magnitude varied across targets, several non-PRC measurements also shifted, and some exclusive populations were small. At 10 kb, the cross-target mean exclusive differences reversed direction for H3K27me3 (*−*0.411), EZH2 (*−*0.485), and SUZ12 (*−*0.584).

Together, these results support using the frozen contact-map representation to retrieve Polycomb-associated contacts through reference examples. At 5 kb, Polycomb guidance favored stronger PRC2-associated enrichment than loop guidance, including among contacts exclusive to each selection. Both populations nevertheless retained PRC2-associated signal, indicating a preference within overlapping contact populations rather than separation into distinct biological classes.

### 2.4 Visually defined retrieval of condensed-domain-like patterns are reproducibly ranked

We next considered patterns that researchers can recognize visually but whose biological identity has not been established. We asked whether reference examples could define such a retrieval query and guide C-Fetch toward patches expressing the requested appearance. Motivated by their resemblance to reported condensed-domain contact-map patterns (Paggi et al. 2026), we used reference patches containing broad, roughly rectangular regions of elevated contact signal extending from the lower-left region toward the patch center. We evaluated four datasets at 5 kb, using references from the other three to guide graph-diffusion retrieval within Mustache proposals without retraining the encoder. To test whether rankings tracked the requested appearance, we separately measured rectangular enrichment using a regional-contrast score that required the lower-left region to exceed both neighboring regions above and to its right (Figure 4A).

**Figure 4:**
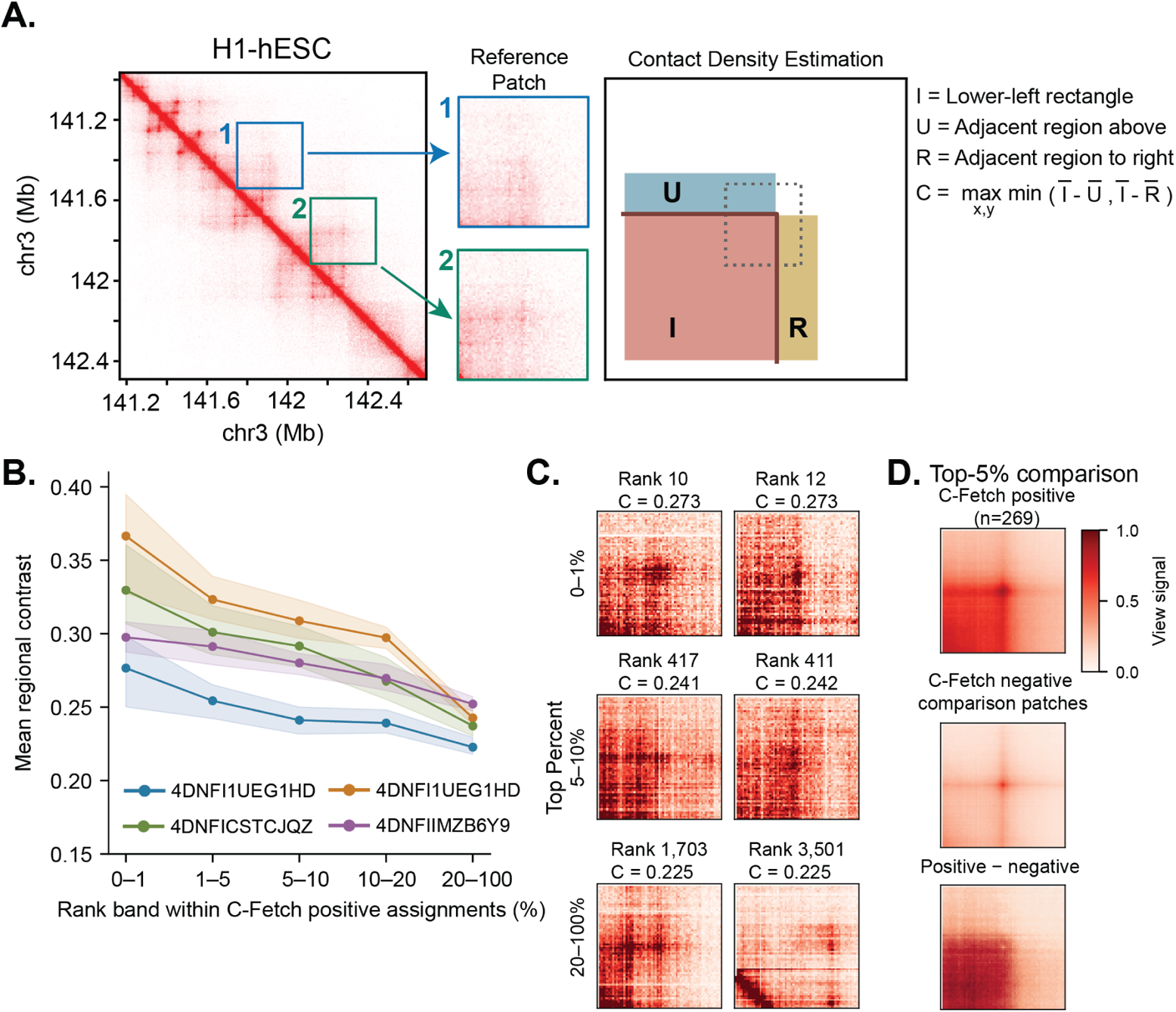
Reference examples guide retrieval of rectangular contact-map patterns across datasets. (A) Reference patches illustrate the queried appearance, and the schematic defines the regional contrast used to assess rectangular enrichment. (B) Higher-ranked C-Fetch-positive assignments showed stronger mean regional contrast within Mustache proposals across four datasets at 5 kb. Datasets are labeled by accession. Shading indicates bootstrap 95% intervals. (C) Examples from 4DNFI1UEG1HD (GM12878) illustrate patches across three rank bands. Rank denotes position among graph-diffusion-positive assignments; *C* denotes regional contrast. Lower rank numbers indicate higher C-Fetch scores, whereas higher *C* indicates stronger rectangular enrichment. (D) The top 5% of positive assignments in 4DNFI1UEG1HD showed stronger lower-left enrichment than C-Fetch-negative comparison patches sampled to resemble their genomic and contact-density characteristics. Positive and negative denote C-Fetch assignments rather than independently validated feature labels.

Higher-ranked C-Fetch-positive patches expressed the requested rectangular appearance more strongly across all four datasets (Figure 4B,C). C-Fetch’s highest-ranked 5% had greater mean regional contrast than the bottom 80% in all four datasets, with means ranging from 0.259 to 0.332 versus 0.223 to 0.253, respectively. Thus, reference examples from other datasets guided C-Fetch toward the requested appearance, and higher-ranked groups expressed that appearance more strongly.

As a supporting comparison, the top 5% also showed greater regional contrast than C-Fetch-negative patches sampled from the same candidate pool to match their genomic location and contact-density characteristics. Positive-minus-negative differences ranged from 0.165 to 0.216 across targets (Figure 4D; Supplemental Figure S4A). Positive and negative here denote graph-diffusion assignments. To assess whether view-scale saturation contributed to this difference, we repeated the comparison while additionally matching the proportion of pixels capped by C-Fetch’s viewing normalization. The differences decreased, remaining positive in 4DNFI1UEG1HD (GM12878) and 4DNFI-IMZB6Y9 (H1-hESC), but becoming near zero in the other two datasets (Supplemental Figure S4). This additional matching retained fewer pairs and did not fully balance saturation, preventing us from isolating its contribution. The positive–negative comparison therefore did not consistently establish greater rectangular enrichment after accounting for saturation, although the separate ranking analysis showed stronger expression of the pattern among higher-ranked positive assignments under the viewing normalization used.

Having retrieved patches by appearance, we explored their compartment and accessibility characteristics to describe their genomic context. Across the four datasets, 52–78% of matched, compartment-annotated top-5% positive assignments had both anchors in A compartments (Supplemental Figure S5). Exploratory ATAC profiles around the fitted rectangular transitions did not show a consistent accessibility peak specific to selected patches (Supplemental Figure S6). Together, these results show that examples can define a visual retrieval query and prioritize patches expressing that appearance within Mustache proposals across datasets. The compartment and accessibility analyses provide exploratory context for the retrieved patches without establishing a shared biological identity or physical condensation.

### 2.5 Loop guided retrieval enriches for focal, cohesin-sensitive contacts

We next considered chromatin loops, an established feature class with a recognizable focal appearance and dedicated detection tools. Here, we asked whether reference examples could help prioritize contacts with stronger focal structure within an existing caller’s proposals. We used loop examples from non-target datasets to guide C-Fetch ranking and selection within Mustache-proposed candidate coordinates. We assessed focal enrichment using aggregate peak analysis (APA), which aligns and averages patches centered on candidate coordinates. The peak-to-lower-left ratio (P2LL) measures the central peak relative to a lower-left background region, with higher values indicating stronger aggregate focal enrichment (Durand et al. 2016; Cao et al. 2020). C-Fetch retained 6.2–14.3% of proposals across four held-out targets (Figure 5A; Supplemental Table S2). We first generated unfiltered APA matrices to preserve the complete proposal set, including candidates near the main diagonal. These aggregates showed compact central peaks for C-Fetch selections, with less diffuse surrounding and main-diagonal signal than the complete Mustache proposal sets (Figure 5B). For quantitative comparison, we applied Juicer’s default 30-bin minimum-distance filter. At 5-kb resolution, this excludes contacts separated by less than approximately 150 kb and prevents strong genomic-distance-dependent diagonal signal from entering the APA window and confounding P2LL with proximity to the diagonal (Durand et al. 2016). Under this standardized off-diagonal comparison, C-Fetch selections showed greater focal enrichment in every map: median P2LL increased from 2.29 for the complete proposals to 3.15 for C-Fetch selections, compared with 0.73 for random coordinates (Figure 5A). To assess whether the ranking prioritized focal structure beyond the final selection threshold, we progressively included lower-ranked candidates. P2LL declined monotonically in every target as these candidates were added (Supplemental Figure S7). Thus, loop reference examples supported both selection of focally enriched subsets from Mustache proposals and a continuous ranking that tracked aggregate focal enrichment.

**Figure 5:**
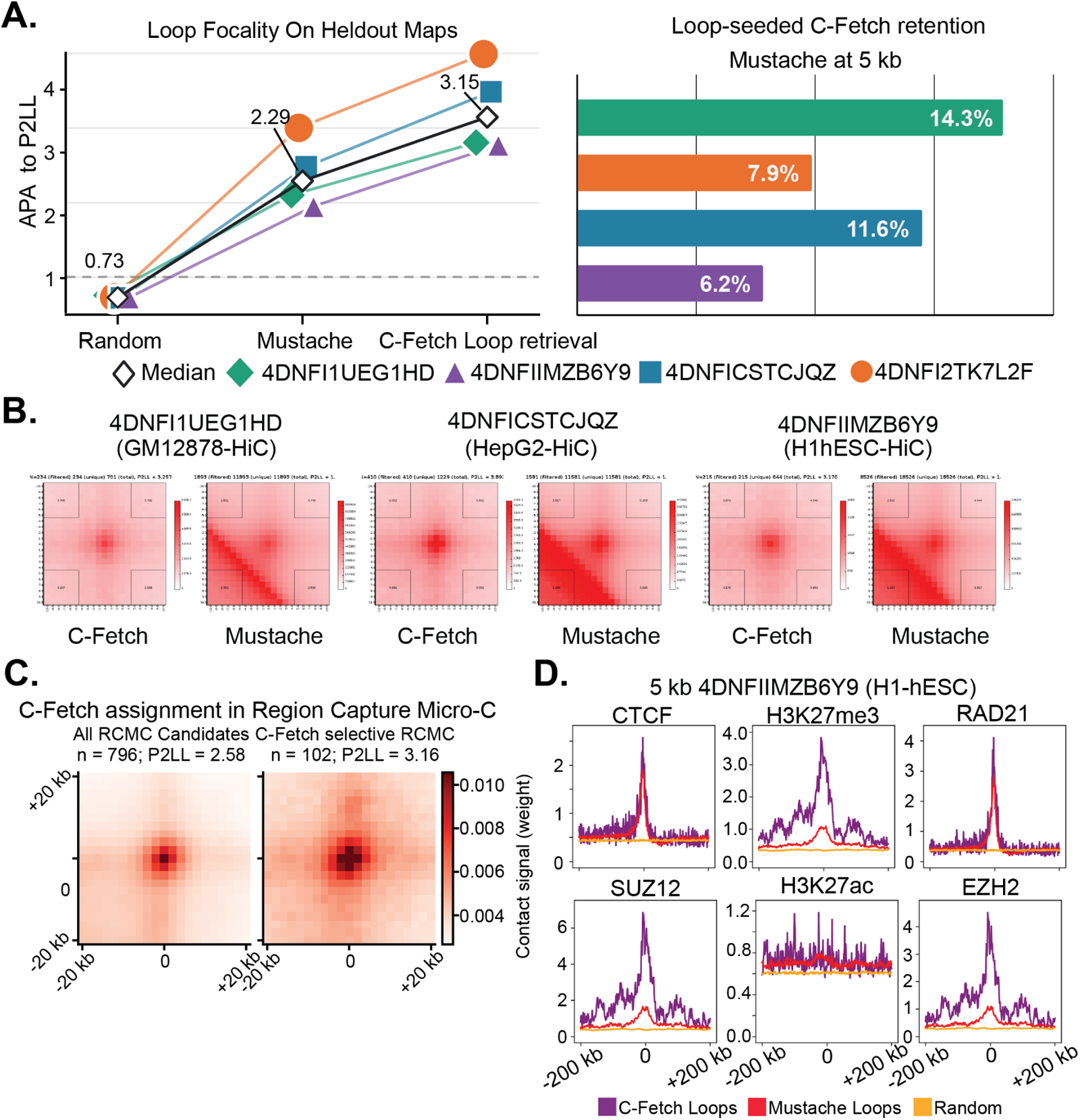
Loop-guided C-Fetch ranking selectively enriches focal off-diagonal contacts within a Mustache-derived proposal set. (A) Quantitative assignment selectivity and focality across four held-out maps. The retained-fraction panel reports the percentage of Mustache proposals assigned by loop-guided C-Fetch. The P2LL panel compares random coordinates, complete Mustache candidate sets, and C-Fetch assignments after excluding contacts within 30 resolution bins of the diagonal. Colored trajectories represent individual maps and open diamonds show medians. Median P2LL was 0.73 for random co-ordinates, 2.29 for Mustache candidates, and 3.15 for C-Fetch assignments; C-Fetch exceeded Mustache in all four maps. (B) Representative 5 kb APA matrices compare loop-guided C-Fetch assignments with the complete Mustache candidate sets in three held-out datasets. C-Fetch aggregates show compact central maxima with less diffuse domain-like and main-diagonal signal. (C) APA of 2 kb GM12878 Region Capture Micro-C candidates. The 102 concordant loop-guided C-Fetch assignments produced greater aggregate focality than all 796 eligible candidates (P2LL 3.16 versus 2.58). Region- and distance-matched resampling is shown in Supplemental Figure S10. (D) H1-hESC ChIP-seq pileups compare C-Fetch loop assignments, Mustache candidates, and random background coordinates for CTCF, RAD21, H3K27me3, EZH2, SUZ12, and H3K27ac. Signal magnitudes are interpreted within each assay and are not compared quantitatively across marks. Because C-Fetch assignments are selected from Mustache proposals, the figure demonstrates selective focality enrichment rather than caller-independent loop recall or feature-level precision.

Because C-Fetch ranks and selects supplied candidates, the caller’s threshold determines which contacts are available for retrieval. We therefore tested whether relaxing that threshold could expand retrieval while preserving earlier selections and focal enrichment. In a separate 5 kb comparison for 4DNFI1UEG1HD, increasing the Mustache significance threshold from 0.01 to 0.1 expanded the candidate pool from 11,882 to 26,108 coordinates. C-Fetch retained 91.3% of its strict-run selections and selected 781 contacts at coordinates absent from the strict pool. Rankings at shared coordinates remained highly concordant (Spearman correlation 0.948; Supplemental Figure S8). C-Fetch selection increased aggregate P2LL within both pools, from 2.10 to 2.65 for the strict pool and from 1.76 to 2.21 for the broader pool. However, the broader pool yielded weaker absolute focality: comparing the top 958 candidates from each run gave P2LL values of 2.64 and 2.43 for the strict and broader pools, respectively (Supplemental Figure S9). Thus, a more permissive candidate list extended reference-guided retrieval while preserving most previous selections, but the focality of the selected population remained dependent on the supplied candidates.

We also evaluated focality within a targeted Region Capture Micro-C candidate pool. Among 796 eligible GM12878 candidates at 2 kb, 102 concordant C-Fetch loop assignments had greater aggregate focality than the complete pool (P2LL 3.16 versus 2.58; Figure 5C). However, the gain was not statistically resolved against region- and distance-matched candidate subsets (empirical P = 0.146; Supplemental Figure S10), indicating that candidate composition could account for the observed difference.

Having shown that C-Fetch selected contacts with stronger aggregate focal enrichment than the complete Mustache proposal sets, we next asked whether this structural prioritization also favored biologically relevant loop behavior. Specifically, we tested whether contacts prioritized in untreated maps showed greater loss of focal enrichment after cohesin depletion than matched subsets of the remaining candidates. Contacts selected from untreated HCT116 maps were evaluated at the same genomic positions after 6 h of Auxin treatment (Rao et al. 2017). Before treatment, C-Fetch-selected contacts had greater aggregate focality than matched subsets of the remaining Mustache candidates at both 5 kb and 10 kb resolution (empirical P = 0.001). Their subsequent focality loss was also greater than the matched-complement distribution at 5 kb (P = 0.002) and 10 kb (P = 0.001; Supplemental Figure S11). Thus, loop-guided prioritization enriched for contacts with cohesin-sensitive focal structure, although the remaining candidates also lost focality after treatment. Epigenomic profiles provided complementary support: H1-hESC loop-guided selections showed narrow central CTCF and RAD21 enrichment, consistent with CTCF/cohesin-associated contacts (Figure 5D) (Rao et al. 2014, 2017). These selections also retained the PRC2-associated enrichment described above, indicating that prioritization of focal structure did not restrict retrieval to a single biological class.

### 2.6 C-Fetch supports transfer across assays and consistent candidate prioritization

Having evaluated biologically and visually defined queries and refinement of existing calls, we next examined how C-Fetch could be reused under different experimental conditions. We tested transfer of the frozen representation to additional contact-map assays, consistency of candidate priorities across sequencing depths and biological replicates, and sensitivity to the number of reference examples.

The frozen model supported reference-guided retrieval on both Region Capture Micro-C and HiChIP maps. On GM12878 RCMC maps, loop and Polycomb queries produced nearly identical selections under an inclusive assignment criterion, whereas requiring agreement between graph diffusion and the calibrated probe yielded smaller selections with more distinct membership between queries (Supplemental Figure S10). On CTCF and SMC1A HiChIP maps, the two queries selected the same candidates at the graph-diffusion threshold but ranked them differently, showing that reference choice could change which contacts were prioritized even when selection membership was unchanged (Supplemental Figure S12). These results support application of the trained representation to additional assays, although gains in aggregate focality over matched candidate subsets were not consistently resolved (Supplemental Figures S10 and S13).

We assessed consistency under reduced sequencing depth and across biological replicates while holding candidate coordinates fixed within each comparison. At one-quarter sequencing depth, correlations between full-depth and downsampled scores ranged from 0.612 to 0.820 across the two reference queries and three assignment procedures, with agreement deteriorating under more severe downsampling (Supplemental Figure S14). Across two GM12878 biological-replicate maps, the combined ranking procedure produced score correlations of 0.731–0.769 when evaluating the same coordinates with the same references and inference settings (Supplemental Figure S15). Thus, C-Fetch retained useful agreement in relative candidate priorities across biological replicates and under moderate coverage reduction, while categorical assignments were more sensitive to sequencing depth.

We also tested sensitivity to reference availability using five random subsampling draws for loopguided retrieval on 11,882 fixed candidates from 4DNFI1UEG1HD at 5 kb. Pools containing only 31 positive and 31 negative examples retained greater aggregate focal enrichment than the complete candidate set, but the number and identity of selected contacts varied substantially across draws (Supplemental Figures S16 and S17). With 310 positive and 308 negative examples, top-decile Jaccard overlap with the full-reference ranking ranged from 0.822 to 0.867 (Supplemental Figure S16). Larger reference pools therefore supported more consistent prioritization, while smaller pools could still retrieve focally enriched contact populations for exploratory analysis in this evaluation.

## 3 Discussion

C-Fetch uses a learned contact-map representation to support retrieval targets specified through reference examples. The applications evaluated here illustrate three uses of this approach: prioritizing contacts with a known biological association within broad focal-contact proposals, retrieving a visually specified pattern without requiring an established biological identity, and refining an existing caller’s proposals for an established feature class. These tasks use the same frozen encoder, with reference examples defining which patterns to prioritize. The practical contribution is therefore a generalizable framework for investigating different contact-map features within supplied candidate pools.

The representation analyses provide complementary support for these comparisons. Neighborhoods reflected annotated feature distinctions, latent proximity tracked patch resemblance, and averaging across clip contexts improved the consistency of similarity rankings (Figure 2). Together, these findings connect the representation to both categorical annotations and graded contact-map appearance. Joint encoding of structurally related patches may contribute to these comparisons, although our experiments did not isolate its contribution from model architecture or companion selection.

Polycomb-associated retrieval illustrates how reference examples can favor a biological association within a candidate population defined by a broader structural criterion. Mustache proposals can include both Polycomb-associated contacts and CTCF/cohesin-associated loops, whose focal appearances overlap. At 5 kb, Polycomb guidance favored stronger H3K27me3, EZH2, and SUZ12 enrichment than loop guidance, including among contacts exclusive to each selection (Figure 3; Supplemental Figure S1). Both populations nevertheless remained PRC2-enriched, indicating a graded biological preference rather than separation into distinct classes. The preference varied with resolution and reversed on average in the 10 kb exclusive-set comparison, showing that its interpretation depends on the scale at which contacts are represented (Supplemental Figure S3). These associations support biological prioritization but do not establish PRC2-dependent contact formation or the mechanisms of individual contacts.

The rectangular-pattern application addresses a different situation: researchers may recognize an appearance before establishing its biological identity or a dedicated detection rule. Reference examples from other datasets guided retrieval toward the requested pattern, and higher-ranked positive assignments expressed stronger rectangular enrichment within Mustache proposals (Figure 4). This supports using examples to investigate a visual feature while its interpretation remains open. The evidence is specific to the evaluated targets and viewing normalization: additional saturation matching weakened the positive–negative comparison, and normalization-independent ranking was not established (Supplemental Figure S4). Compartment and accessibility profiles provided exploratory genomic context but did not establish physical condensation or a shared biological identity (Supplemental Figures S5 and S6).

For an established feature class with dedicated detection tools, loop-guided retrieval demonstrates how examples can refine prioritization within an existing caller’s proposals. Loop-guided retrieval shows how reference examples can help researchers select focally enriched contacts from an existing caller’s proposals. C-Fetch selections had greater aggregate focality than the complete Mustache proposal sets, and focality declined as lower-ranked candidates were included, linking the continuous ranking to the strength of the retrieved contact pattern (Figure 5A,B; Supplemental Figure S7). Contacts prioritized before cohesin depletion also showed greater initial focality and a greater subsequent focality loss than matched subsets of the remaining candidates, providing independent biological support for the prioritization (Supplemental Figure S11) (Rao et al. 2017). Together, these findings support using C-Fetch to focus follow-up on contacts with pronounced, perturbation-sensitive focal structure. Because the measurements describe aggregate populations, they do not establish individual-contact precision or imply that excluded proposals are false positives. Relaxing the Mustache threshold expanded retrieval while preserving most earlier selections and substantial ranking agreement, although the broader selections had lower absolute focality (Supplemental Figures S8 and S9). Thus, the supplied candidate pool determines both the contacts available for retrieval and the structural enrichment that reference-guided selection can achieve.

Reuse of the frozen representation extended to additional assays, although the evidence for transfer differed across settings. On Region Capture Micro-C maps, it retrieved focal contacts, although additional focality beyond region- and distance-matched candidate subsets was not statistically resolved (Figure 5C; Supplemental Figure S10). On CTCF and SMC1A HiChIP maps, changing references altered candidate rankings even when categorical selections were identical, while focality gains over matched controls were not consistent across queries (Supplemental Figures S12 and S13). With candidate coordinates fixed, rankings also retained useful agreement across biological replicates and under moderate downsampling, but agreement deteriorated with more severe coverage reduction (Supplemental Figures S14 and S15). These findings support use of continuous rankings for comparative prioritization across the tested settings, while categorical membership and the strength of retrieved enrichment remain sensitive to assay and coverage.

Reference availability places a further practical condition on reuse. Small reference collections retained aggregate focal enrichment in the loop experiment but produced more variable selections, whereas larger collections supported more consistent prioritization (Supplemental Figures S16 and S17). Their benefit was improved consistency rather than a monotonic increase in focality or assignment count. These results support exploratory queries with limited annotations while showing that the reference requirements for stable retrieval must be evaluated for each application.

Together, these applications show how one contact-map representation can support biologically motivated retrieval, exploration of visually specified patterns, and refinement of established feature calls. Reference examples define the target class, candidate coordinates are assigned and ranked against the class, and structural or biological measurements provide the evidence needed to interpret each retrieval. Existing callers and the supplied coordinate sampler offer ways to usefully constrain candidate coordinates; further queries and sampling strategies will establish how broadly the approach extends. C-Fetch provides a generalized framework for this exploration without requiring a new model or tool for each feature.

## 4 Methods

### 4.1 Reference data

All datasets were obtained from public repositories; accessions, experimental sources, datasets or cell lines, assay types, and primary references are provided in Supplemental Tables S5–S9. Training data comprised seven Hi-C and two Micro-C datasets, from which Mustache identified approximately 300,000 loop coordinates at 2, 5, and 10 kb resolution in hg38. Within each chromosome and source dataset, we sampled additional non-overlapping patches matched to the called coordinates by genomic span and distance from the diagonal, yielding 20.6 million Hi-C and 8.0 million Micro-C patches in total, sampled equally from each data source at 2, 5, and 10 kb (Supplemental Tables S5 and S6). Evaluation candidates were generated using Mustache and random patch sampling from three held-out Hi-C datasets and one held-out Micro-C dataset (Supplemental Tables S7 and S8). Additional datasets included CTCF and SMC1A HiChIP, GM12878 Region Capture Micro-C, untreated and auxin-treated HCT116 Hi-C, and ChIP-seq for epigenomic analyses (Supplemental Table S9).

### 4.2 Contact-map image patches

Contact-map patches were extracted from observed contact values without matrix normalization. Candidate-centered windows spanned approximately 320 kb along each matrix axis: 324 kb at 2 kb and 320 kb at 5 kb and 10 kb.

For both training and retrieval, patches were max-scaled after excluding the main diagonal identified using Otsu thresholding and Hough detection, falling back to the full-patch maximum when diagonal masking was unavailable. Values were clipped to [0, 1], non-finite values replaced with zero, and patches resized to 128 *×* 128 pixels by bilinear interpolation.

### 4.3 Non-temporal clip construction

To construct non-temporal clips, we grouped contact-map patches by similarity in sliding-window Fourier features. Within working sets of 8,000 patches, k-means clustering grouped patches with related spatial-frequency patterns. Within clusters containing up to 512 patches, a greedy nearest-neighbor walk ordered patches by Euclidean distance between their Fourier features; larger clusters were ordered by projection onto the leading principal component. The ordered patches were stored in eight-frame records, with leftover patches pooled into additional records. During training, five frames were sampled without replacement from each record to provide overlapping contexts. The clip axis represents groups of patches rather than time; 3D convolutions operate across this axis without a next-frame or causal objective (Tran et al. 2015; Tong et al. 2022; WanTeam et al. 2025).

### 4.4 Modeling rationale

C-Fetch uses an autoencoding bottleneck to preserve contact-map information needed for local comparison while reducing nuisance variation (Bengio et al. 2013; Kingma and Welling 2014). Reconstruction alone does not guarantee a useful retrieval space, and regularized autoencoders need not follow a simple prior closely to support smooth local representations (Vincent et al. 2008; Rifai et al. 2011; Ghosh et al. 2020).

### 4.5 Manual annotation of reference feature sets

We manually annotated candidate-centered patches using C-Fetch’s input genomic extents for four datasets (4DNFI1UEG1HD, 4DNFI2TK7L2F, 4DNFICSTCJQZ, and 4DNFIIMZB6Y9) at 2, 5, and 10 kb resolution, with approximately 400 examples per class and condition. Loop references contained an approximately centered, discrete focal contact without a visible main diagonal, repeated or grid-like puncta, prominent stripes, or domain-corner structure. Feature-negative references lacked a discernible central feature but could contain off-center signal. Polycomb-associated references contained at least four focal contacts in an approximately square or rectangular array. These appearance-based criteria defined operational classes rather than establishing molecular identity (Rao et al. 2014; Kraft et al. 2022). The latent-space benchmark used exactly 400 loop and 400 negative examples per dataset–resolution condition, totaling 9,600 examples.

For the visually defined dense-chromatin query, we annotated approximately 100 patches per dataset at 5 kb resolution. These contained concentrated rectangular enrichment extending from the lower-left region toward, and potentially across, the patch center. This class specified a visual retrieval target without assuming an established biological identity.

### 4.6 Weakly KL-regularized autoencoders

The 2D autoencoder follows the Wan-VAE design, adapted to normalized single-channel 128 *×* 128 contact-map patches, with encoder and decoder widths of 128 and 256 channels, respectively (Wan-Team et al. 2025). The encoder retained a diagonal Gaussian posterior parameterization with weak KL regularization, while reconstruction used the posterior mean without latent sampling (Kingma and Welling 2014). Training used L1 reconstruction loss, SSIM in place of LPIPS (Wang et al. 2004), and KL regularization toward a standard normal prior, weighted by 3, 0.05, and 3 *×* 10*^−^*^6^, respectively.

The 3D model was initialized from the 2D checkpoint by placing learned weights in the center slice of each 3D kernel and using symmetric padding along the non-temporal clip axis (Tran et al. 2015; WanTeam et al. 2025). It was trained for an additional 100,000 steps on five-patch inputs (Section 4.3) using the same objective over clip reconstructions.

### 4.7 Reconstruction evaluation

We evaluated the 2D and 3D autoencoders on held-out normalized Hi-C patches and non-temporal clips, respectively, using MSE, MAE, PSNR, and SSIM. Reconstructions and targets were evaluated on the same normalized intensity scale; clip SSIM was averaged across frames. We also measured FID (Heusel et al. 2017) and an adapted FVD using pretrained 3D ResNet-18 embeddings (Unterthiner et al. 2018; Hara et al. 2018), interpreting these as relative feature-distribution measures rather than Hi-C-specific perceptual scores.

### 4.8 Latent-space evaluation

We evaluated posterior-mean embeddings from the 135M and 650M models on the 9,600-example benchmark defined in Section 4.5, using five-patch inputs. Neighborhood purity was the mean same-label fraction among the *k* = 25 nearest neighbors under RMS-L2 distance. Results were calculated within each dataset–resolution condition and averaged with equal condition weights; pooled analyses recomputed neighborhoods across the combined examples. For radius sweeps, coverage was the fraction of queries with at least five neighbors within the specified radius, and purity was calculated among covered queries. Radii were treated as model-specific rather than calibrated across independently trained representations.

Complementary analyses assessed standardized cosine similarity, same-class versus cross-class ranking, latent norms, purity lift, and nearest-neighbor margins. We visualized embeddings using PCA followed by t-SNE (van der Maaten and Hinton 2008) and assessed local decoder sensitivity using finite-difference Jacobian estimates and second directional differences. Detailed diagnostic and visualization settings are provided in Supplemental Methods.

### 4.9 Reference-guided feature assignment

The frozen encoder mapped labeled reference patches and target candidates into a shared representation. For each target dataset and resolution, references originating from that dataset were excluded, although homologous genomic loci could occur across datasets. Comparisons between loop and Poly-comb guidance changed only the reference examples, holding the encoder, candidate coordinates, latent transformation, and assignment settings fixed.

#### Inference clips and embeddings

Inference used four-patch clips, distinct from the five-patch training inputs. Reference clips were constructed within each label, dataset, resolution, and coordinate source. Candidates were grouped into structural pools using multiband short-window Fourier-transform features. Each candidate appeared as the designated subject in 16 clips, each containing three related context patches selected by a deterministic balanced sampler. Flattened clip posterior means were centered and RMS-normalized within each vector. Clips were scored individually before evidence was aggregated for each candidate; embeddings were not averaged before assignment. Detailed pooling and sampling settings are provided in Supplemental Methods.

#### Distance-decayed conformal assignment

Reference source rows were partitioned before clip construction into disjoint proper-reference and calibration sets, reserving 25% for calibration. Class-conditional nonconformity scores were defined by the seventh-nearest-neighbor RMS-L2 distance to proper references (Shafer and Vovk 2008). Calibration scores yielded conformal values, which were weighted by distance decay and a smooth calibration-boundary penalty. Candidate evidence combined subject-clip and context-partner evidence with weights of 0.75 and 0.25, respectively. Acceptance required sufficient conformal support, evidence, and agreement across clips. These scores were used as calibrated evidence without claiming cross-dataset coverage guarantees.

#### Graph-diffusion assignment

We constructed weighted nearest-neighbor graphs of reference and candidate clip embeddings within each structural pool, adding direct connections to references from both classes. Reference-label evidence was propagated using a row-normalized transition matrix with restart and clamped reference labels (Zhou et al. 2003). Assignments required sufficient propagated class probability and direct connectivity to reference nodes; near-tied class probabilities were marked ambiguous.

#### Calibrated linear-probe assignment

A class-balanced linear softmax classifier was fitted to frozen embeddings standardized using the proper-reference partition (Alain and Bengio 2016). Class-conditional calibration combined classifier probabilities with distances to reference-class centroids. Assignments required sufficient calibrated support and classifier probability, limited centroid distance, and agreement across clips. Otherwise accepted clips were additionally rejected when their leading evidence fell below the 90th percentile across the query population. Complete scoring equations, optimization settings, and clip- and candidate-level acceptance thresholds for all three procedures are provided in Supplemental Methods.

#### Assignments and consensus ranking

Each procedure returned categorical assignments and continuous scores. Categorical consensus required agreement between at least two procedures; otherwise, candidates were designated NoConsensus. Inclusive selections required positive support from any procedure, whereas graph–probe concordant selections required support from both graph diffusion and the linear probe.

For continuous consensus ranking, procedure-specific scores were converted to percentile ranks within each target, reference query, and coordinate set, assigning tied scores their mean rank. The three percentile ranks were averaged, providing an ordering even for candidates without a positive categorical assignment.

#### Candidate-coordinate scope

Mustache coordinates contributed to unlabeled training-patch sampling and defined the principal evaluation candidate sets, but caller labels were not encoder-training targets. The analyses therefore assess retrieval within supplied proposals rather than caller-independent genome-wide recall.

### 4.10 Assay-transfer, perturbation, and robustness analyses

All analyses used the frozen encoder; reference examples and inference settings were held fixed within comparisons except when reference-pool size was explicitly varied. Detailed protocols and parameter settings are provided in Supplemental Methods.

#### Candidate-pool expansion

We compared loop-guided retrieval from Mustache candidate sets generated at significance thresholds of 0.01 and 0.1 for 4DNFI1UEG1HD at 5 kb resolution. We measured selection retention, ranking agreement among shared coordinates, and aggregate focal enrichment, including comparisons of equal-sized selections. Focality comparisons were descriptive and were not matched for genomic distance.

#### Assay transfer

We evaluated GM12878 Region Capture Micro-C at 2, 5, and 10 kb resolution using non-GM12878 references, comparing loop- and Polycomb-guided selections under inclusive and graph–probe concordant criteria. For HG00731 CTCF and SMC1A HiChIP at 10 kb, we compared graph-diffusion rankings and selection overlap under the two reference queries. HiChIP focal enrichment was assessed against equal-sized, genomic-distance-matched random subsets of eligible candidates.

#### Reference availability and sequencing depth

Reference-pool sensitivity was evaluated on a fixed GM12878 candidate set at 5 kb using nested, source-stratified positive and negative reference subsets across five random draws. Sequencing-depth sensitivity was evaluated at fixed full-depth candidate coordinates using progressively downsampled maps. We measured assignment recovery, Spearman score correlation, and Jaccard overlap of selected sets relative to the corresponding full-reference or full-depth baseline; reference subsampling also included aggregate focal-enrichment comparisons.

#### Biological-replicate stability

Mustache candidate sets from two GM12878 biological replicates were each evaluated on both maps at 5 kb using external references. Agreement was measured by Spearman score correlation and top-20% Jaccard overlap, using category-aware priority rankings for individual procedures and the continuous rank-fused consensus.

#### Auxin perturbation

We compared untreated and 6-h auxin-treated HCT116 maps from GSE104333 at 5 and 10 kb. Candidates were called from the untreated map, and contacts supported by both graph diffusion and the calibrated probe were selected once in that condition and evaluated unchanged in both maps. Focal enrichment and its treatment-associated decrease were compared with the remaining candidates using paired candidate bootstraps and equal-sized, chromosome- and genomic-distance-matched null subsets.

### 4.11 Aggregate peak analysis

We measured focal enrichment using KR-normalized aggregate peak analysis (APA) in Juicer Tools (Durand et al. 2016), separately from model-input normalization. For the primary 5 kb analysis, APA used 21*×*21 matrices and a minimum anchor separation of 30 bins. P2LL was the central pixel divided by the mean of the 6*×*6 lower-left background block. HiChIP aggregates instead used balanced cooler matrices. Analysis-specific settings and null comparisons are provided in Supplemental Methods.

### 4.12 Epigenomic enrichment and selection overlap

Epigenomic signal was averaged within *±*25 kb of each anchor, then across anchors so that each contact contributed one observation. Enrichment was the log_2_ ratio of mean signal in a selected set to that in the shared random-patch background for the corresponding dataset and resolution, with a fixed pseudocount. Polycomb-versus-loop differential enrichment was the difference between these enrichment values.

Selections were matched by exact coordinates, allowing anchor swaps, and partitioned into shared and query-exclusive contacts. We estimated 95% percentile intervals using 5,000 contact-level boot-strap replicates (Efron and Tibshirani 1993). Comparisons of complete and exclusive selections used a paired joint bootstrap preserving their shared membership. The primary aggregate analysis combined four target datasets and both window sizes at 5 kb resolution.

### 4.13 Evaluation of visually defined rectangular patterns

We evaluated graph-diffusion retrieval from Mustache candidates across four datasets at 5 kb, using references from the other three datasets. Positive assignments were grouped by score percentile; percentiles refer to the positive population rather than all candidates.

Rectangular enrichment was measured independently of retrieval scores on viewing-normalized patches. For each candidate rectangle, contrast was the smaller of its mean-intensity differences from adjacent strips above and to its right; the maximum contrast across tested boundaries was retained. Rank-band means were summarized with 95% percentile intervals from 400 chromosome-pair boot-strap resamples. Displayed retrieved examples were selected near each band’s median contrast.

Positive assignments were compared with graph-negative patches matched by chromosome pair, genomic distance, intensity, and nonzero-pixel fraction, allowing negative-patch reuse. Sensitivity analyses additionally matched the fraction of pixels clipped during normalization and reported pair retention and residual imbalance. These analyses assessed sensitivity to comparison-set matching rather than changes to C-Fetch inference. Boundary-search settings, matching procedures, and aggregate-image construction are detailed in Supplemental Methods.

### 4.14 Software availability

The C-Fetch inference software, including model loading, database ingestion, clip construction, latent encoding, the three assignment procedures, rank-fused consensus generation, paper configurations, and example command-line workflows, is available at https://github.com/seanpatrickmoran/chromatin-fetch. Code for extracting contact-map patches into fp32 matrices and cataloging them in SQLite databases is available in the rustrainer repository at https://github.com/seanpatrickmoran/rustrainer; patch extraction can be accessed through strainer.py. The accompanying coordinate sampler provides statistical sampling, optional distance-decay weighting, and full tiling to prepare candidate coordinates for this extraction and inference workflow.

### 4.15 AI-assisted writing and coding

During preparation of this work, the authors used ChatGPT for LaTeX formatting, manuscript editing, and coding assistance. The authors reviewed and edited the resulting content and take full responsibility for the publication.

## 5 Data access

### Annotated C-Fetch reference data

The annotated contact-map patches and reference examples used for C-Fetch retrieval and evaluation are available through Hugging Face at https://huggingface.com/datasets/signaldetectionweary/cfetch-annotations. The release contains cell-line-, resolution-, and label-specific SQLite databases, a tabular annotation manifest without serialized image arrays, database schemas, label definitions, source-dataset manifests, and file checksums. The source contact maps from which these patches were extracted remain available from the repositories and accessions listed in Supplemental Tables S5–S9.

### Model checkpoints

The trained C-Fetch representation-model checkpoints, model configurations, preprocessing metadata, and usage documentation are available through Hugging Face at https://huggingface.co/signaldetectionweary/chromatin-fetch-135M and https://huggingface.co/signaldetectionweary/chromatin-fetch-650M. The base 650M-parameter 3D checkpoint was used for the primary retrieval and assignment analyses reported in this study.

## 7 Acknowledgments

The study was supported by the National Human Genome Research Institute (R35HG011279) and by the National Heart, Lung, and Blood Institute (R01HL170115). The authors would like to acknowledge Linghua Jiang and Kiaxuan Jin for their invaluable feedback on the paper.

## Author contributions

Sean Moran (SM) and Jie Liu (JL) contributed to study design and interpretation of the results. SM developed the methodology and software, curated and annotated the datasets, trained the models, performed the computational and statistical analyses, prepared the figures, and drafted the manuscript. JL supervised the research and contributed to manuscript revision. Both authors reviewed and approved the final manuscript.

